# Development of binding and activity inhibition assays for the antibiotic resistance-associated protein PhoQ

**DOI:** 10.64898/2026.06.15.732377

**Authors:** Hannah G. Addis, Dawson L. Blankenship, Erin E. Carlson

## Abstract

Antimicrobial resistant infections present a growing threat to public health and were associated with or directly caused 6 million deaths globally in 2021. This huge death toll highlights the need for novel strategies to address AMR infections. Interfering with the regulation of resistance mechanisms could provide an alternative approach to treat drug-resistant infections. PhoQ, a sensor histidine kinase ubiquitous amongst gram-negative bacteria, regulates several virulence factors, as well as resistance to outer membrane-targeting antibiotics, making it an attractive target for adjuvant therapy development. However, the identification of potent small molecule inhibitors is limited by the assays available for *in vitro* assessment of binding and activity inhibition in PhoQ. Thus, we sought to investigate the use of a fluorescence-based assay to evaluate enzymatic activity, as well as a thermal shift assay to assess inhibitor-protein binding in PhoQ. Together, these newly implemented protocols are valuable contributors to the toolbox of methods available for the development of PhoQ-targeted inhibitors to block this major contributor to antimicrobial resistance.

## Introduction

Antimicrobial resistant (AMR) infections are predicted to cause or contribute to more than 10 million death by 2050^1^. The rapid rise in AMR coupled with the slowed introduction of new antibiotics into the clinic make the need for novel strategies to address these infections urgent^2–4^. A promising approach aims to intercept bacterial virulence and resistance mechanisms to resensitize infections to once-resistant antibiotics^5^. By inhibiting these behaviors, adjuvants disarm the pathogen and enable existing antibiotics to clear resistant pathogens.

Two-component systems (TCS) often regulate virulence and resistance behaviors making them important targets for therapeutic development^6, 7^. TCSs, typically consisting of a membrane bound histidine kinase (HK) and its cognate response regulator (RR), are signal transduction pathways that respond to external stimuli to elicit a cellular response, such as activation of antibiotic resistance mechanisms (**Figure 1A**)^8^. Environmental signals interact with the signaling domain of the HK and trigger ATP-mediated autophosphorylation of a conserved histidine residue^9^. This phosphoryl group is transferred to the RR, which often regulates gene expression^9^. Within the HKs, ATP is bound by a Bergerat fold, a series of structural homology boxes in the catalytic ATP-binding domain (CA domain), making this site attractive for broad-spectrum inhibition^10^. Indeed, we anticipate that molecules targeting the CA domain could be used to disarm multiple virulence and resistance pathways in numerous species^11^. Herein, we report the validation and use of two assays to further enable the discovery and optimization of PhoQ inhibitors that target its highly conserved CA domain.

**Figure 1.**
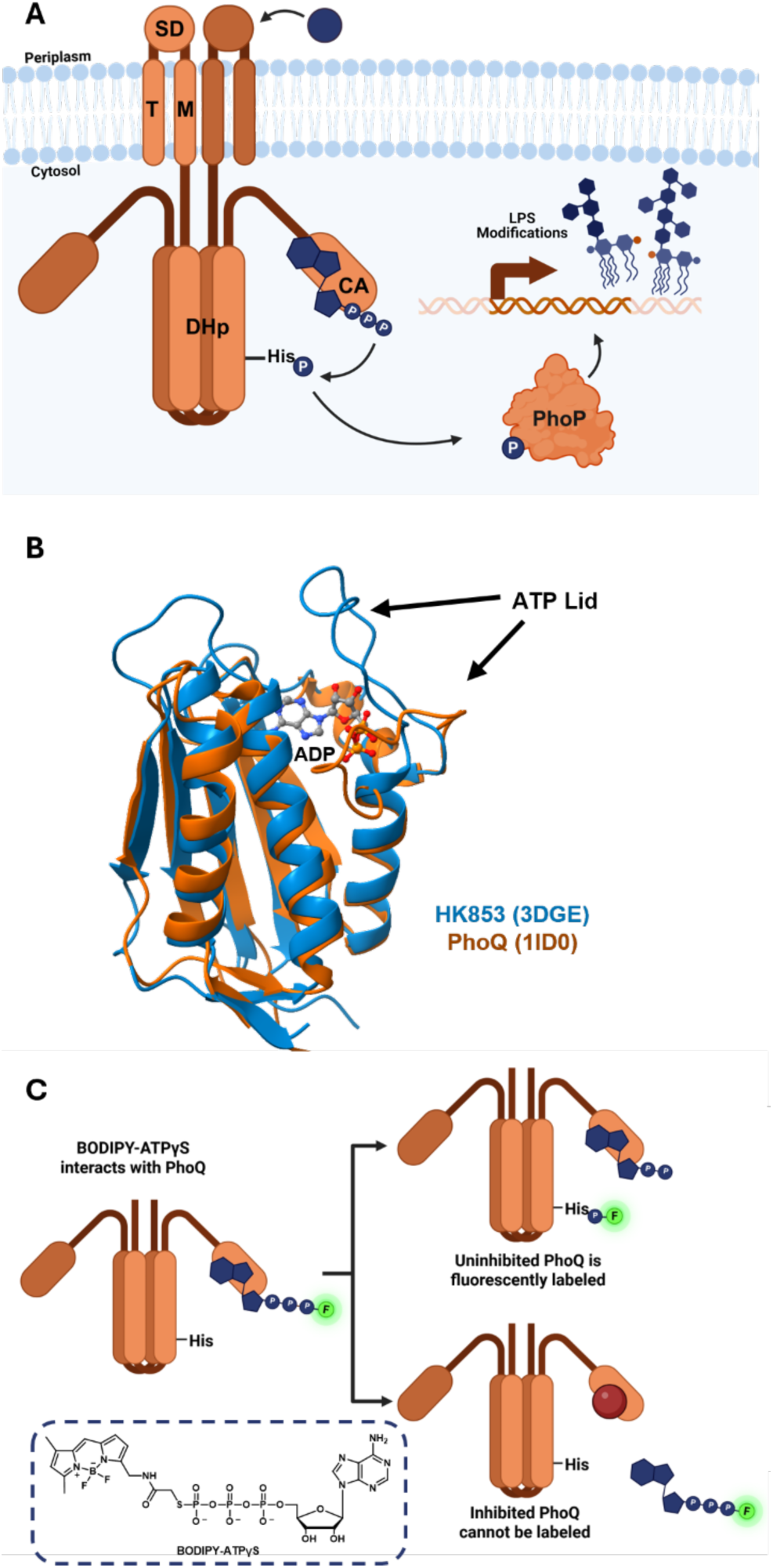
Mechanism of the PhoP/Q TCS in regulating polymyxin resistance and activity-based labeling of PhoQ. (A) Signals interacting with the SD of PhoQ promote autophosphorylation of a conserved histidine via ATP bound to the CA domain. The phosphoryl group is transferred to the PhoP response regulator, which can then modulate various phenotypes, including LPS modifications that confer polymyxin resistance. SD = signaling domain, TM = transmembrane domain, DHp = dimerization histidine phosphorylation domain, CA = catalytic ATP-binding domain. (B) Alignment of HK853 (PDB 3DGE) and PhoQ (PDB 1ID0) CA domains, including the flexible ATP-lid, in ChimeraX demonstrating conserved structural homology between proteins. (C) Fluorescent probe BODIPY-FL-ATPγS (B-ATPγS) interacts with uninhibited PhoQ to fluorescently label the conserved histidine of catalytically active protein. Inhibition of PhoQ at the CA domain prevents labeling, enabling readout of PhoQ activity.

We have previously focused much of our HK inhibitor discovery work on HK853, a model protein from *Thermatoga maritima*. HK853 is a thermostable protein with a well-characterized structure and mechanism, readily enabling screening and optimization studies.^12, 13^ However, it originates from a marine bacterium with no ties to infection or known links to virulence^14–16^. These drawbacks highlight the need for use of an infection-relevant protein to assess small molecule binding and activity inhibition.

PhoP/Q (RR/HK) is a Mg^2+^ sensing TCS found in multiple strains of pathogenic bacteria, including *Salmonella enterica* and *Enterobacter cloacae*^17^. It regulates virulence- and resistance-associated processes, such as modification of lipid A, a component of outer membrane lipopolysaccharide (LPS), to confer resistance to polymyxin antibiotics^18^ (**Figure 1A**). Moreover, it has also been structurally and mechanistically characterized and exhibits structural homology with the CA domain of HK853 (**Figure 1B**)^17, 19–21^. In addition, HK inhibitors sharing a 2-aminobenzothiazole scaffold discovered by our lab have been implicated in PhoQ inhibition and potentiation of polymyxin resistance in *S. enterica* and *E. cloacae*^22, 23^. However, we had not yet obtained evidence of direct interaction of these molecules with PhoQ. Herein, we report the optimization and application of multiple inhibitor assays with PhoQ, through use of our previously reported activity-based assay^13^, as well as differential scanning fluorimetry. We evaluated a small library of 2-aminobenzothiazole HK inhibitors that resensitize gram negative bacteria to polymyxins, implying engagement of PhoQ, for their ability to bind and inhibit this protein *in vitro*. Two of the tested molecules bind to PhoQ and only one inhibits autophosphorylation activity, with a much lower potency than in HK853, indicating that despite the highly conserved character of their ATP-binding domain, selectivity between HKs can be achieved via inhibition at this pocket.

## Materials and Methods

Additional information regarding protocols and materials can be found in the Supporting Information.

### Protein Expression, Purification, and Preparation

The codon optimized sequence for *T. maritima* HK853_Cyt_ (residues 235–492) and *S. enterica* PhoQ_Cyt_ (residues 332-487) was inserted into the pET28a(+) plasmid by Twist Bioscience. Protein was expressed using *E. coli* BL21 (DE3) cells in Miller LB and induced with 0.2 mM IPTG at 20°C overnight. Cells were harvested by centrifugation (10,000 x *g* for 30 min at 4 °C) and resuspended in 40 mL Lysis Buffer (25 mM Tris, 500 mM NaCl, 10 % glycerol, 2 mM BME, 20 units DNAse, pH 8, one Roche cOmplete, EDTA-free protease inhibitor cocktail tablet). The cell resuspension, kept on ice, was lysed by probe sonication (15 sec on, 30 sec off, 15 min total on time, 30% amplitude). Cell lysates were centrifuged at 13,000 x *g* for 45 min at 4 °C and supernatant was filtered using a 0.22 μm filter. Filtered lysate was loaded onto a HisTrap crude column (Cytiva). The column was washed with buffer A (50 mM Tris, pH 7.8, 500 mM NaCl, 10% glycerol, 2 mM BME) and the bound proteins were eluted using a linear gradient of buffer B (50 mM Tris, pH 8, 500 mM NaCl, 10 % glycerol, 2 mM BME, 500 mM imidazole). Fractions containing the desired protein were exchanged into storage buffer (10 mM Tris, 0.1 mM EDTA, 12% glycerol, 2 mM BME, pH 7.6), concentrated using Amicon centrifugal filters and stored at −80 °C. The final purity was greater than 90% by SDS-PAGE. On the day of an experiment, purified protein was exchanged into reaction buffer (50 mM Tris, 200 mM KCl, 5 mM MgCl_2_, pH 7.8, 0.1% Triton X-100) using 10 kDa molecular weight cut off filters and protein concentration was determined via BCA microplate assay.

### Activity Assay Optimization

To determine the ideal time and temperature conditions, 23.75 μL of 1 μM PhoQ_Cyt_ was aliquoted into microcentrifuge tubes for each time point within a temperature. B-ATPgS probe (1.25 μL of 50 μM) was added to each aliquot and was vortexed and briefly centrifuged. Samples were covered in aluminum foil and incubated at their respective temperatures. At each time point, the reaction was quenched with 8.6 μL 4X SDS loading dye (200 mM Tris, 400 mM DTT, 8% SDS, 0.2% bromophenol blue, 40% glycerol, pH 6.8) and stored at 4 °C during the remaining time points. Once all reactions were quenched, samples were run on 10% Tris polyacrylamide gels 180V for 1h. Gels were washed three times with Milli-Q water before being read on a Typhoon 9210 Gel Scanner on the BODIPY channel. Total protein levels were obtained using Coomassie staining. Gels were uniformly processed in ImageJ to remove background, increase contrast, and measure band integrated density. The same protocol, with reactions run at 25 °C for 3 h, was used to study the effects of a protein concentration curve on labeling. To test assay tolerance to DMSO, the same protocol with 4 μM PhoQ_Cyt_, 20 μM B-ATPgS, and various concentrations of DMSO was used. A t-test comparing each DMSO condition to an untreated control in Graphpad Prism was used to determine any statistical significance, p-value < 0.05.

### Kinetics

Kinetics evaluation in PhoQ_Cyt_ and HK853_Cyt_ was performed using methods as previously described.^24^

### Activity Assay Dose-Response Curves and IC_50_ Value Determination

Dose-response curves and IC_50_ value determinations were executed as previously reported for HK853_Cyt_, with changes to the optimized parameters (4 μM PhoQ_Cyt_, 20 μM B-ATPγS, three-hour incubation time).

### DSF

To test binding via DSF, 96-well PCR plates were set up to test compound dilution curves in triplicate. Twelve-compound dilution curves were prepared in DMSO and 1.25 μL of each was added to designated wells to give a final concentration range 0 – 300 μM . Protein was diluted in reaction buffer containing no Triton X-100 to a concentration of 25 μM, and 5 μL of protein stock was added to designated wells, with reaction buffer adding up to 20 μL total volume for a final protein concentration of 5 μM. Controls including each compound concentration with dye alone were included to ensure no background effects. Plates were centrifuged at 200 rpm for 5 min and incubated at room temperature for 25 min. SYPRO Orange dye was diluted to a 25X stock using the same reaction buffer, and 5 μL were added to designated wells for a final concentration of 5X. Plates were sealed with microseal film and centrifuged once more before being read on a Bio-Rad CFX Duet Real-Time PCR System. The melt curve spanned 25 – 95 °C, with temperatures increasing 0.5 °C/30 sec and measurements taken at every increase using the FRET channel. Melting temperatures were calculated by dRFU using DSFworld.^25^

## Results and Discussion

### Optimization of a fluorescence-based activity assay in PhoQ

Evaluating the activity of HKs is limited by the unstable nature of the phosphohistidine bond, which hydrolyzes too quickly for traditional kinase assays to be useful.^26^ Instead, most HK activity assays, including studies analyzing the activity of PhoQ, use radiolabeled ATP and SDS-PAGE autoradiography or phosphate affinity electrophoresis using Phos-tag SDS-PAGE to quantify activity.^15, 20, 27, 28^ Autoradiography has the drawback of requiring the use of radiation and the implementation of specific safety protocols, making the procedure tedious and time-consuming to perform while also exposing researchers to harmful radiation. Instead, we used a previously developed a fluorescence-based assay that employs BODIPY-FL-ATPγS thioester (B-ATPγS), a probe that displays a sulfur on the gamma phosphate of ATP, which yields a stabilized thiophosphohistidine bond. Formation of a covalent bond between the protein and fluorophore when a HK is catalytically active can be detected following SDS-PAGE by fluorescent gel analysis. Initial application of this assay has been with a cytosolic construct of HK853 (HK853_Cyt_; **Figure 1C**).^13^ This protein, along with the *S. enterica* PhoQ construct reported herein, (PhoQ_Cyt_),^20^ lacks the sensory and transmembrane regions and possesses only the dimerization histidine phosphorylation (DHp) domain and CA domain appended to an N-terminal ^6^His tag for purification. These constructs are constitutively active, enabling the identification of activity inhibitors targeting the CA domain.^13, 20^ However, our assay had not previously been adapted to the study of infection-relevant HKs, such as PhoQ.

To determine the optimal conditions for PhoQ_cyt_ reaction with our activity-based probe, B-ATPγS, they were co-incubated at temperatures ranging from 4 °C to 37 °C for up to six hours (**Figure 2A**). Labeling of the active protein, determined by integrated density of SDS-PAGE gel bands under the BODIPY channel, directly correlated to the time of the reaction enabling us to quantify PhoQ autophosphorylation kinetics.^24^ While no PhoQ_cyt_ labeling occurred at any tested time point at 4 °C, enzyme activity was detected between 25-37 °C with similar levels of PhoQ_cyt_ labeling. As such, we selected the lowest incubation temperature and time that yielded reproducible signal intensity, 3 hours at 25 °C, for the assessment of PhoQ_cyt_ activity.

Using these conditions, the lowest concentration of protein to yield an intense signal was determined (**Figure 2B**). The level of labeling increased as the amount of PhoQ_cyt_ increased, as expected, and 4 μM protein was selected as this minimized protein consumption with maximal labeling.

**Figure 2.**
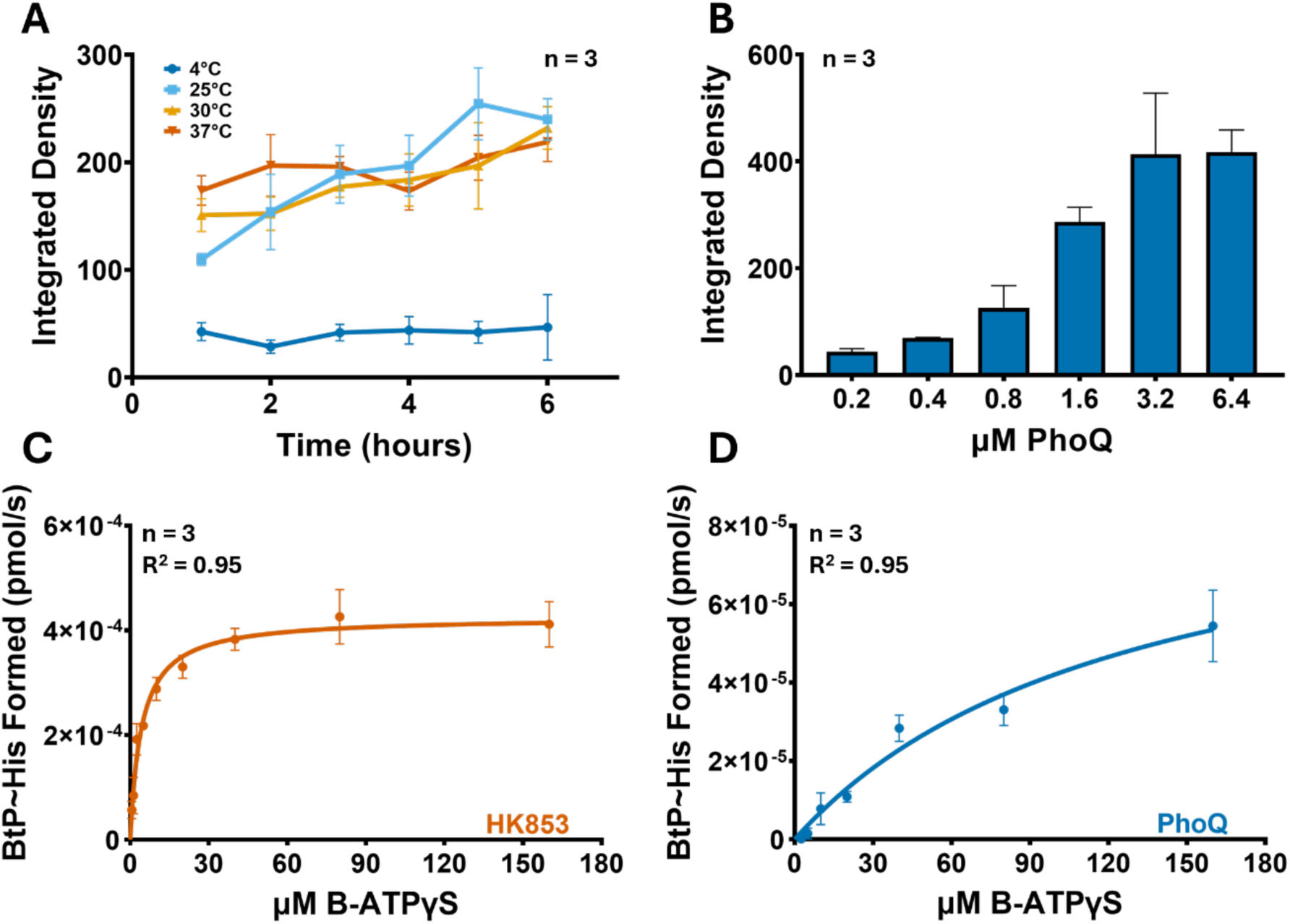
Optimization of fluorescence-based HK activity assay in PhoQ_Cyt_. (A) Integrated band density as a measure of 1 μM PhoQ_Cyt_ labeling with 2.5 μM B-ATPγS over time and at various temperatures. (B) Integrated band density over a variety of PhoQ concentrations at 25 °C and 3 h incubation. (C) B-ATPγS concentration-dependent autophosphorylation of histidine (BtP∼His) in HK853_Cyt_. Reactions were fit to the Michaelis-Menten function to calculate the kinetic parameters summarized in **Table 1**. (D) B-ATPγS concentration-dependent autophosphorylation of histidine (BtP∼His) in PhoQ_Cyt_. Reactions were fit to the Michaelis-Menten function to calculate the kinetic parameters summarized in **Table 1**.

### PhoQ_cyt_ labeling is slower and less efficient than HK853

The increased reaction time needed for adequate PhoQ_Cyt_ labeling from one hour for HK853 to three hours indicates a slower reaction time or decreased affinity for the probe, prompting further kinetic studies. Reaction kinetics were determined as previously described^24^. Kinetics experiments were also run on HK853_Cyt_ as a positive control and the data were compared to previously reported values for HK853 and VicK, a TCS implicated in cell wall synthesis regulation from *Streptococcus pneumoniae*, to gain insights into PhoQ_Cyt_’s relative reaction efficiency (**Table 1**)^24^.

The obtained kinetic parameters of HK853_Cyt_ are comparable to the previously published values from our group.^24^ As anticipated, PhoQ_Cyt_ is labeled much more slowly and has decreased affinity for B-ATPγS, making its turnover efficiency with this probe similar to VicK. While labeling efficiency is relatively low, it is sufficient to use this assay for future studies. Finally, these data enabled us to determine the probe concentration at half-maximal binding to use moving forward (40 μM). Together, these optimization experiments establish conditions under which PhoQ_Cyt_ activity and inhibition can be evaluated with B-ATPγS.

**Table 1.** Comparison of kinetic parameters for HKs in reaction with B-ATPγS. Parameters determined by fitting data from Figure 2C and 2D to a Michaelis-Menten curve in GraphPad Prism. Shaded rows indicate previously reported data.^24^

| PROTEIN | K<sub>m</sub> (μM) | k<sub>cat</sub> (s<sup>-1</sup>) | V<sub>max</sub> (pmol/s) | k<sub>cat</sub>/K<sub>m</sub> (M<sup>-1</sup>s<sup>-1</sup>) |
| --- | --- | --- | --- | --- |
| HK853<sub>Cyt</sub> | 6.27 ± 0.86 | (1.92 ± 0.0730) x 10<sup>-5</sup> | (3.84 ± 0.15) x 10<sup>-4</sup> | 3.06 |
| VicK<sub>Cyt</sub> | 147 ± 21.0 | (5.93 ± 1.02) x 10<sup>-6</sup> | (3.19 ± 2.04) x 10<sup>-4</sup> | 0.0400 |
| HK853<sub>Cyt</sub> | 4.31 ± 0.98 | (3.68 ± 0.104) x 10<sup>-5</sup> | (4.25 ± 2.42) x 10<sup>-4</sup> | 6.45 |
| PhoQ<sub>Cyt</sub> | 129 ± 51.6 | (2.42 ± 0.87) x 10<sup>-6</sup> | (9.67 ± 7.39) x 10<sup>-5</sup> | 0.0190 |

### Adenosine-based molecules prevent fluorescent labeling of PhoQ_cyt_

With conditions for probe labeling of PhoQ_Cyt_ established, known binders of the PhoQ CA domain were investigated to confirm that they are competitive with B-ATPγS as expected, resulting in decreased protein labeling. Additon of ATP (substrate) or ADP (product), as well as a non-hydrolyzable version of ATP (AMP-PNP) resulted in a dose-dependent decrease in PhoQ_Cyt_ labeling (**Figure 3**). From these data, IC_50_ values for each compound were calculated (**Table 3**), which are consistent with reported K_d_ values for adenosine-based compounds with PhoQ^29^. Next, we evaluated a small library of HK inhibitors, previously identified by our group^30, 31^, against PhoQ to provide comparative data for this resistance-relevant HK. The selected 2-aminobenzothiazole compounds showed good inhibition (low micromolar potencies) against HK853 *in vitro* (**Table 2**). R7, which replaces the 2-amino group with a methyl, eliminates inhibitory activity in HK853_Cyt_ and was included as a negative control^31^.

We were particularly interested in learning more about the potential of R1 and R2, both containing a 2-aminobenzothiazole scaffold, to interact with PhoQ given their activity as adjuvants in polymyxin sensitivity assays performed in *S. enterica* and *E. cloacae*.^22, 23^ We reported that R1 caused a sixteen fold decrease in MIC of colistin and polymyxin B against polymyxin-resistant *S. enterica* in a plate-based checkerboard assay.^22^ We also found that R1 (100 μM) and R2 (500 μM) potentiate colistin-mediated killing of *E. cloacae*.^23^ Resistance to this class of antibiotics is partially conferred by PhoQ activation.^17, 21^

**Figure 3.**
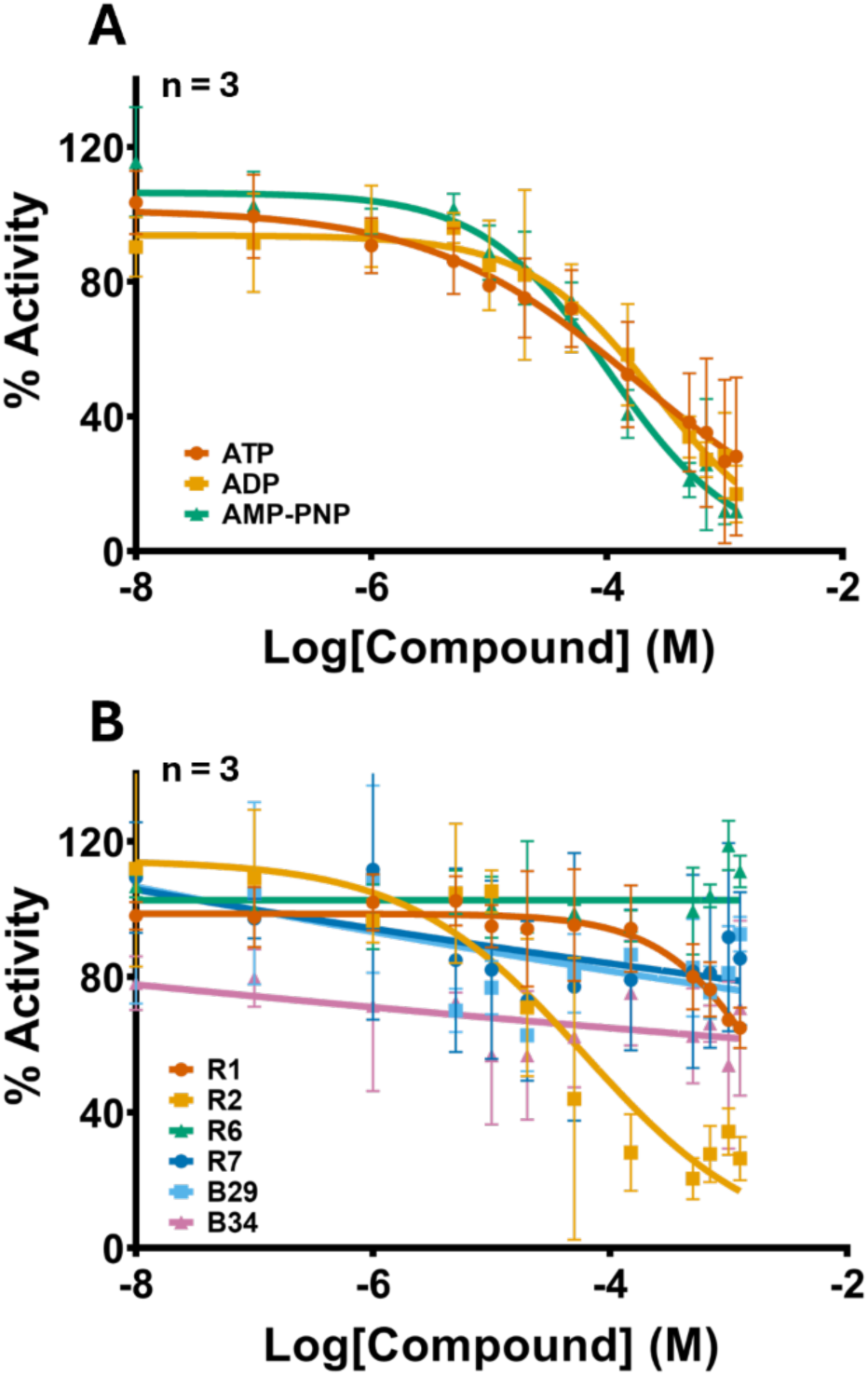
Evaluation of PhoQ activity inhibition with known HK binders. (A) Dose-response curves of adenosine-based ligands in PhoQ_Cyt_. (B) Dose-response curves of a small 2-aminobenzothiazole library for PhoQ_Cyt_ activity inhibition. IC_50_ values included in **Table 3**.

Of the six 2-aminobenzothiazoles tested with PhoQ, only R2 inhibited its activity with an IC_50_ of 60.4 μM, 95% CI [17.04 – 165.1] (**Table 3**). This represents a 30-fold decrease in inhibitor potency compared to HK853. This result was surprising given the ability of R2 to potentiate colistin killing indicating that future structure-activity-relationship studies that aim to improve potency in PhoQ_Cyt_ could provide a more potent adjuvant for polymyxin treatment. The lack of inhibition of the remaining molecules in PhoQ_Cyt_ compared to HK853_Cyt_ indicate that the “dimer”-like structure of R2 may be essential for adequate PhoQ_Cyt_ binding. Discrepancies in the inhibition of all tested molecules also underscores a nuance in designing pan-inhibitors of HKs that the highly conserved homology of the CA domain alone is insufficient for translating small molecule binders between proteins, further highlighting the need for use of infection-relevant HKs for *in vitro* studies.

**Table 2.** Structures of a small library of 2-aminobenzothiazole compounds and their IC_50_ values for HK853_Cyt_ inhibitory activity.

| Compound | R<sub>A</sub> | R<sub>B</sub> | HK853 IC<sub>50</sub><br>(μM) | 95% CI |
| --- | --- | --- | --- | --- |
| R1 | NH<sub>2</sub> | OCF<sub>3</sub> | 7.15 | 6.73 – 7.59 |
| R2 | NH<sub>2</sub> |  | 1.21 | 1.03 – 1.42 |
| R6 | NH<sub>2</sub> | CF<sub>3</sub> | 15.1 | 9.98 – 22.7 |
| R7 | CH<sub>3</sub> | NH<sub>2</sub> | >1250 | - |
| B29 | NH<sub>2</sub> |  | 3.10 | 1.34 – 6.47 |
| B34 | NH<sub>2</sub> |  | 0.420 | 0.190 – 0.820 |

### Differential scanning fluorimetry to evaluate binding to PhoQ_cyt_

Differential scanning fluorimetry (DSF), a common method for evaluating small molecule binding, has no established conditions for use in PhoQ. DSF is a straightforward and high-throughput method for determining small molecule binding via a shift in protein melting temperature, a strategy often used in drug development,^32, 33^ making this assay an ideal complement to the above-described activity assay.

**Figure 4.**
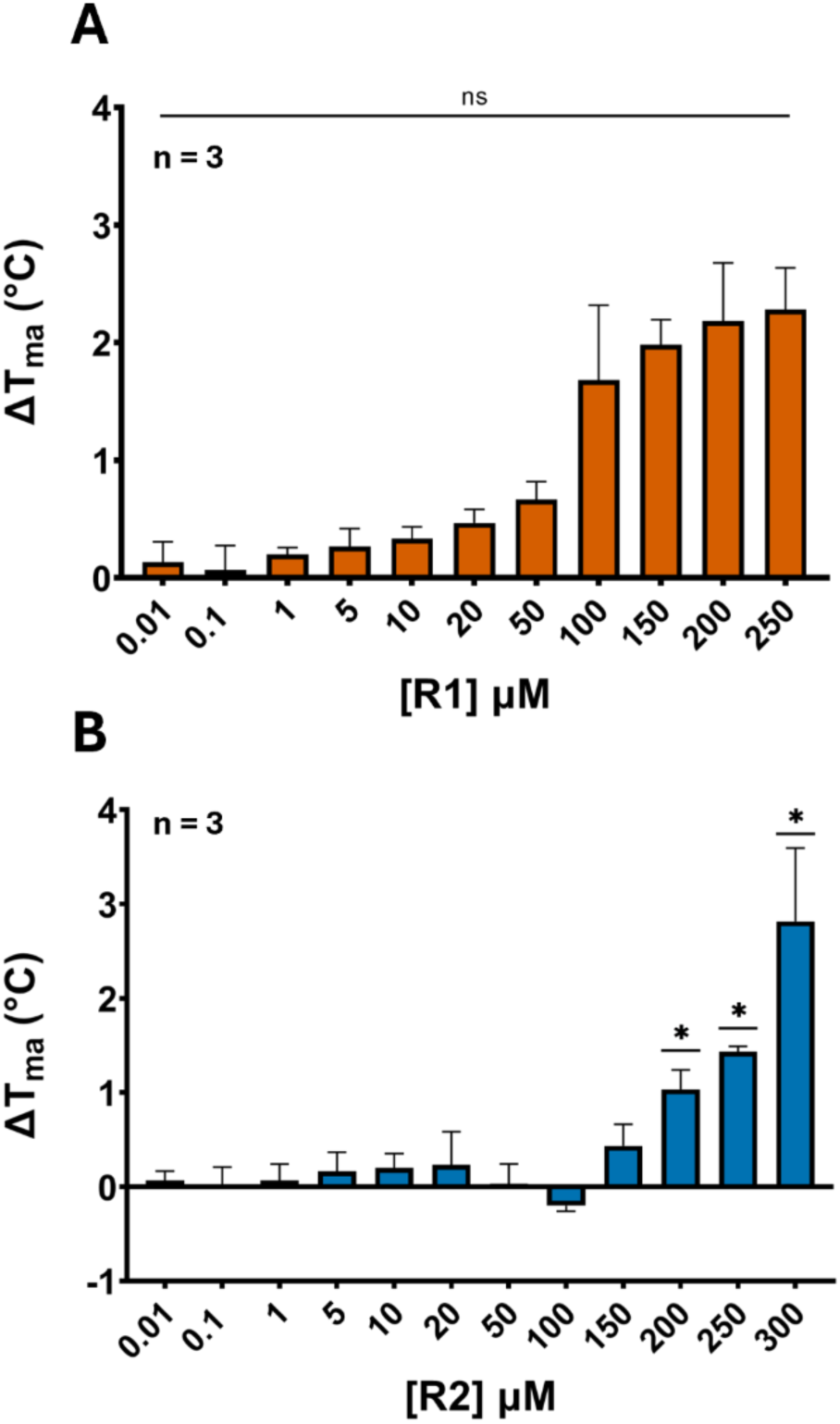
Differential scanning fluorimetry to evaluate small molecule binding to PhoQ_Cyt_. (A) Change in melting temperature (T_m_) of R1 over a range of compound concentrations. (B) Change in T_m_ of R2 over a range of compound concentrations. Significance determined using unpaired t-test comparisons between the T_m_ of samples compared to the DMSO control. P-value significance indicated using asterisks (* ≤ 0.01).

Following condition optimization, the melting temperature of PhoQ_Cyt_, both untreated and with DMSO added was calculated^34^. Next, we added varying concentrations of each molecule to test for the dose-dependent change in melting temperature (ΔT_m_) that is indicative of protein stabilization/destabilization upon small molecule binding (**Figure S5A**; **Table 3**). ADP was used as a positive control, in which an increase of 3 °C in melting temperature was observed at 300 μM (**Figure 4B**). We found that of the 2-aminobenzothiazoles, only R2 exhibits a significant dose-dependent increase in T_m_ indicating PhoQ binding (**Figure 4C and 4D**). R2 was the only molecule to both bind PhoQ_Cyt_ and inhibit protein activity, validating previous data showing potentiation with polymyxins. R1, even though it had no impact on PhoQ_Cyt_ activity and no statistically significant change in T_m_ from the DMSO control, exhibits a dose-dependent increase. This implies that it could bind at higher concentrations and likely impacts polymyxin resistance in *S. enterica* through a pathways that does not involve direct interaction with PhoQ.

**Table 3.** Summary of IC_50_ and ΔT_m_ values with PhoQ. Shaded compounds are adenosine-based controls. NT = not tested.

| Compound | IC<sub>50</sub> (μM) | 95% CI | $\Delta T_m$ at 250 μM (°C) |
| --- | --- | --- | --- |
| ATP | 177 | 64.7 – 375 | 3.20 |
| ADP | 255 | 157 – 393 | NT |
| AMP-PNP | 104 | 70.9 – 149 | NT |
| R1 | >1250 | - | 2.00 |
| R2 | 60.4 | 17.0 – 165 | 2.80 |
| R6 | >1250 | - | 0.70 |
| R7 | >1250 | - | 1.30 |
| B29 | >1250 | - | -0.100 |
| B34 | >1250 | - | 0.900 |

## Conclusions

Herein, we report the optimization and use of two *in vitro* assays for the study of small molecule inhibitors with PhoQ – one to evaluate compound binding and the other to assess inhibition of enzymatic activity. While we have utilized a fluorescent activity assay, which uses B-ATPγS as an activity-based probe, in multiple investigations focused on the model HK, HK853, it had not been applied to the study of another HK. Careful optimization enabled us to translate this assay to PhoQ and provides important comparative data to those previously obtained with this model protein. To complement these studies, we established DSF with PhoQ to investigate small molecule-protein binding interactions in this antibiotic resistance-related HK.

Discrepancies between the 2-aminobenzothiazole library’s potency in PhoQ vs HK853 highlight the importance of developing HK inhibitor assays in pathogenesis-related proteins. More studies will be needed to understand how the structural differences, like the flexible ATP-lid region, between the HK853 and PhoQ CA domain contribute to this gap in activity and how molecule design can be adapted to improve potency against PhoQ. Further structure-activity relationship studies using these PhoQ assays with an expanded library of 2-aminobenzothiazole compounds would expand our understanding of small molecule inhibition of HKs to block bacterial virulence and resensitize resistant bacteria to polymyxins.

## ACCESSION CODES

HK853: UniProtKB Q9WZV7

PhoQ: UniProtKB D0ZV89

VicK: UniProtKB A0A0H2ZNH9

## ASSOCIATED CONTENT

## AUTHOR INFORMATION

### Corresponding Author

*Erin E. Carlson

### Author Contributions

H.G.A conceived of this project, performed most of the experiments and wrote the paper draft; D.L.B. expressed and purified PhoQ_Cyt_ protein and helped perform experiments shown in Figure 2; E.E.C. contributed to the design and completion of this project and manuscript editing.

## ACKNOWLEDGMENT

This work was supported by NIH MIRA R35GM153306.

## TOC FIGURE

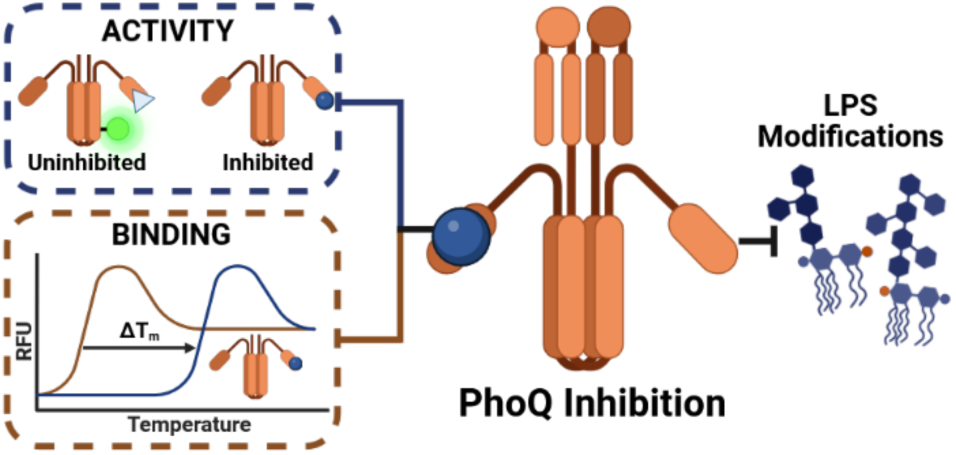

Optimization of novel activity and binding assays for investigating interactions with PhoQ expands the current chemical biology toolbox and allows for the evaluation of small molecule inhibitors of histidine kinases as potential therapeutics for antimicrobial resistant bacteria.

