## Supporting Information for "Development of binding and activity inhibition assays for the antibiotic resistance-associated protein PhoQ"

**Supporting Tables and Figures**

**Figure S1.** SDS-PAGE gels for optimization of time, temperature, protein concentration, and DMSO concentration in the $PhoQ_{Cyt}$ activity assay

**Figure S2.** Standard curves and SDS-PAGE gels used to quantify $PhoQ_{Cyt}$ and $HK853_{Cyt}$ labeling in kinetics experiments

**Figure S3**. SDS-PAGE gels for compounds tested in $PhoQ_{Cyt}$ activity assay

**Figure S4**. Change in melting temperature for each compound in $PhoQ_{Cyt}$

**Figure S5**. RFU curves of tested compounds in DSF with $PhoQ_{Cyt}$.

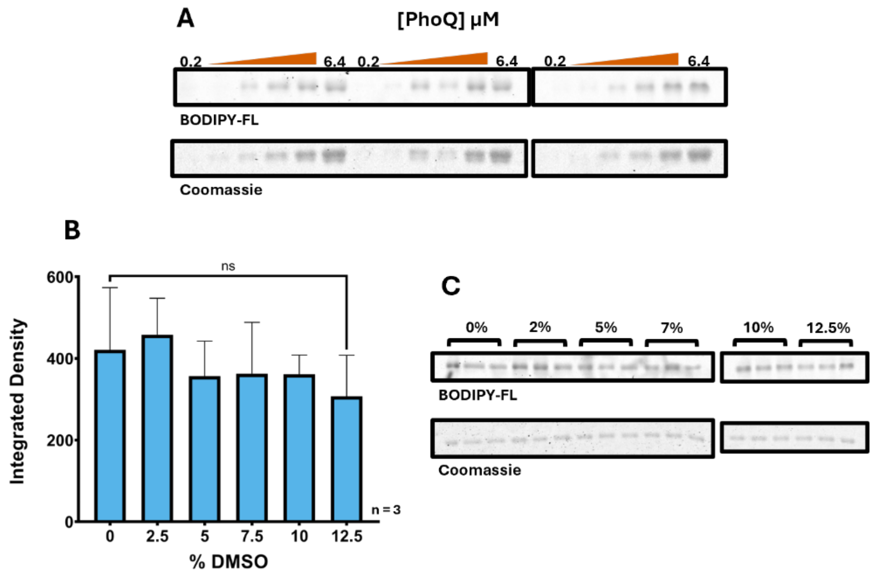


**Figure S1.** SDS-PAGE gels for optimization of (A) protein concentration and (C) DMSO concentration in the $PhoQ_{Cyt}$ activity assay. (B) Integrated densities of DMSO treatments were graphed and found to not differ significantly compared to the 0% control by an unpaired t-test.

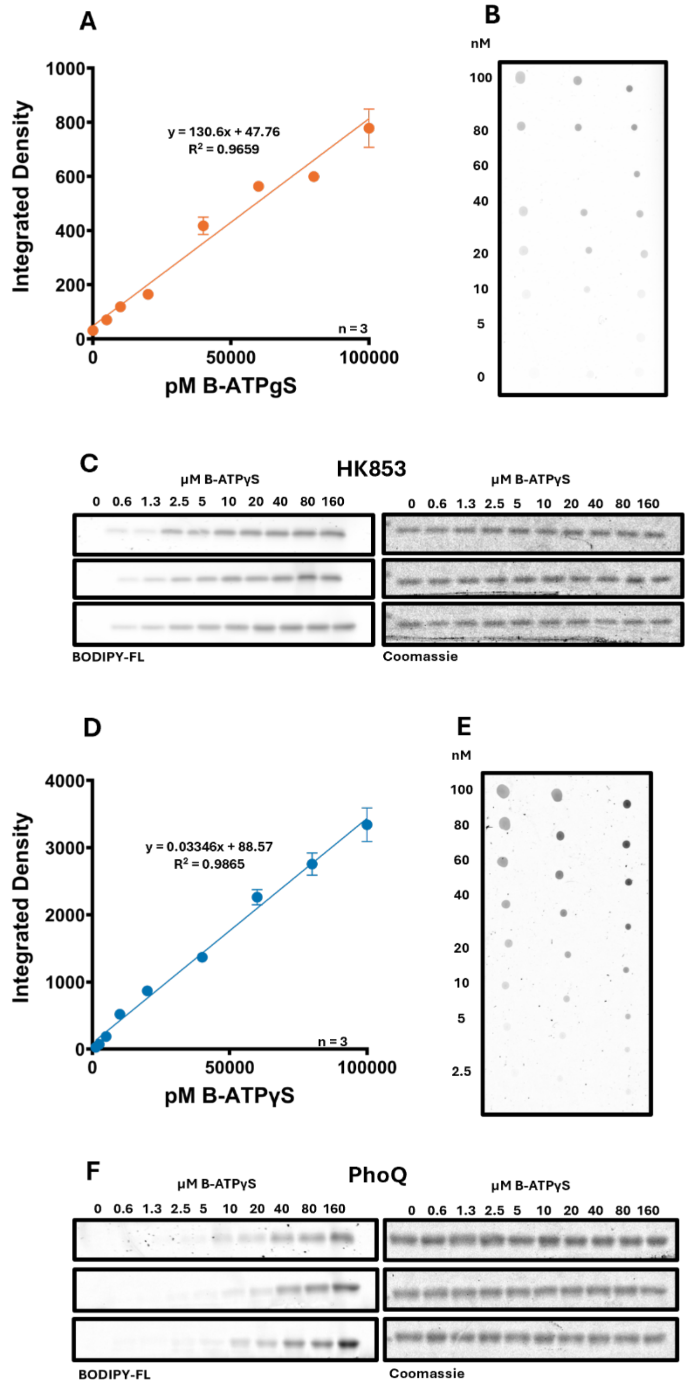


**Figure S2.** Standard curves and SDS-PAGE gels used to quantify $PhoQ_{Cyt}$ and $HK853_{Cyt}$ labeling in kinetics experiments. (A) Integrated densities of a standard curve of probe alone fitted to a linear trendline for labeling quantification in $HK853_{Cyt}$ kinetics experiments. (B) Probe standard curve spotted on a plastic sheet protector for use in $HK853_{Cyt}$ kinetics experiments. (C) SDS-PAGE gels from $HK853_{Cyt}$ kinetics experiments scanned in both BODIPY-FL and Coomassie channels. (D)

Integrated densities of a standard curve of probe alone fitted to a linear trendline for labeling quantification in $PhoQ_{Cyt}$ kinetics experiments. (E) Probe standard curve spotted on a plastic sheet protector for use in $PhoQ_{Cyt}$ kinetics experiments. (F) SDS-PAGE gels from $PhoQ_{Cyt}$ kinetics experiments scanned in both BODIPY-FL and Coomassie channels.

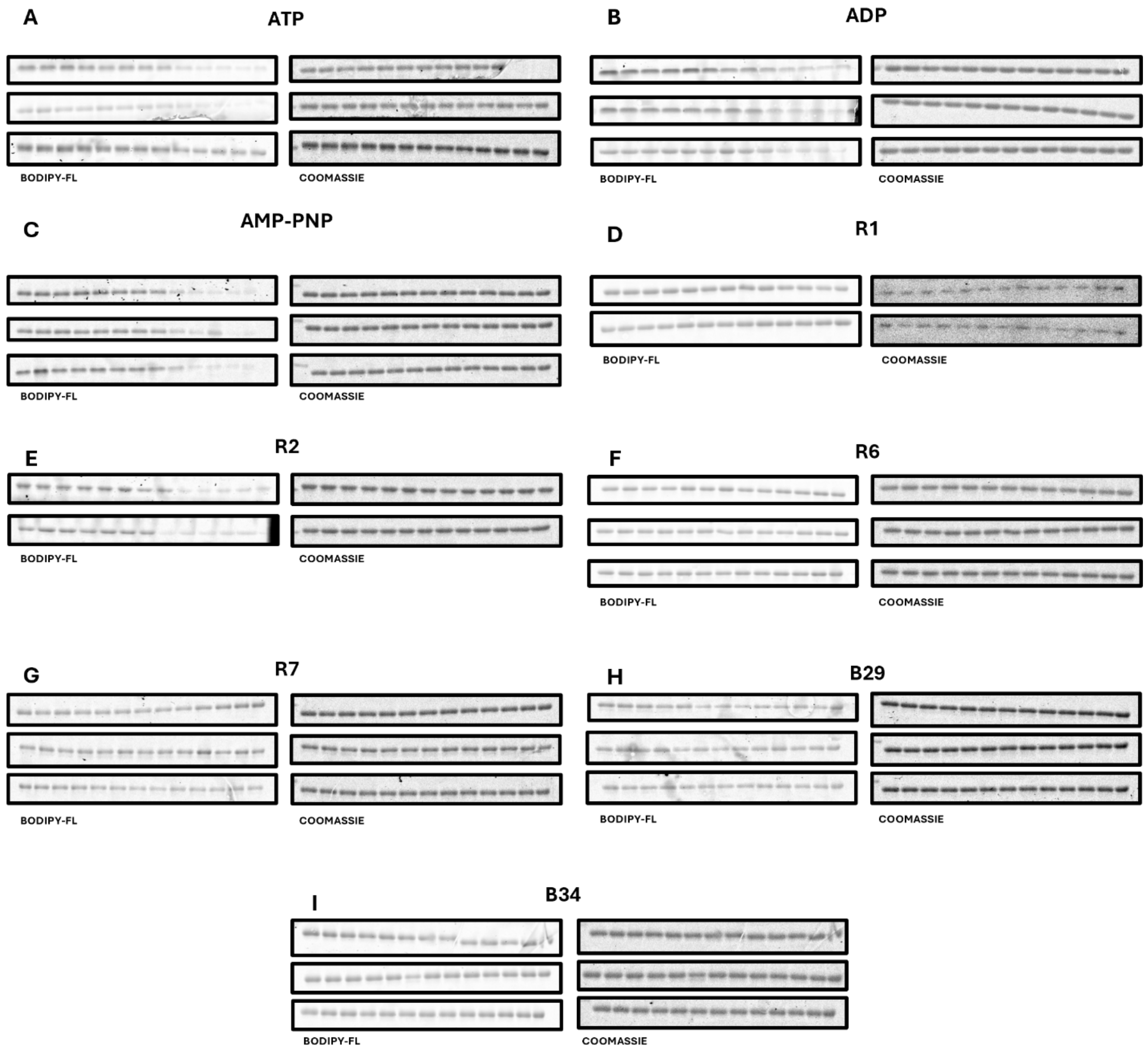


**Figure S3**. SDS-PAGE gels for compounds tested in $PhoQ_{Cyt}$ activity assay. (A-I) All compounds scanned in both BODIPY-Fl and Coomassie channels.

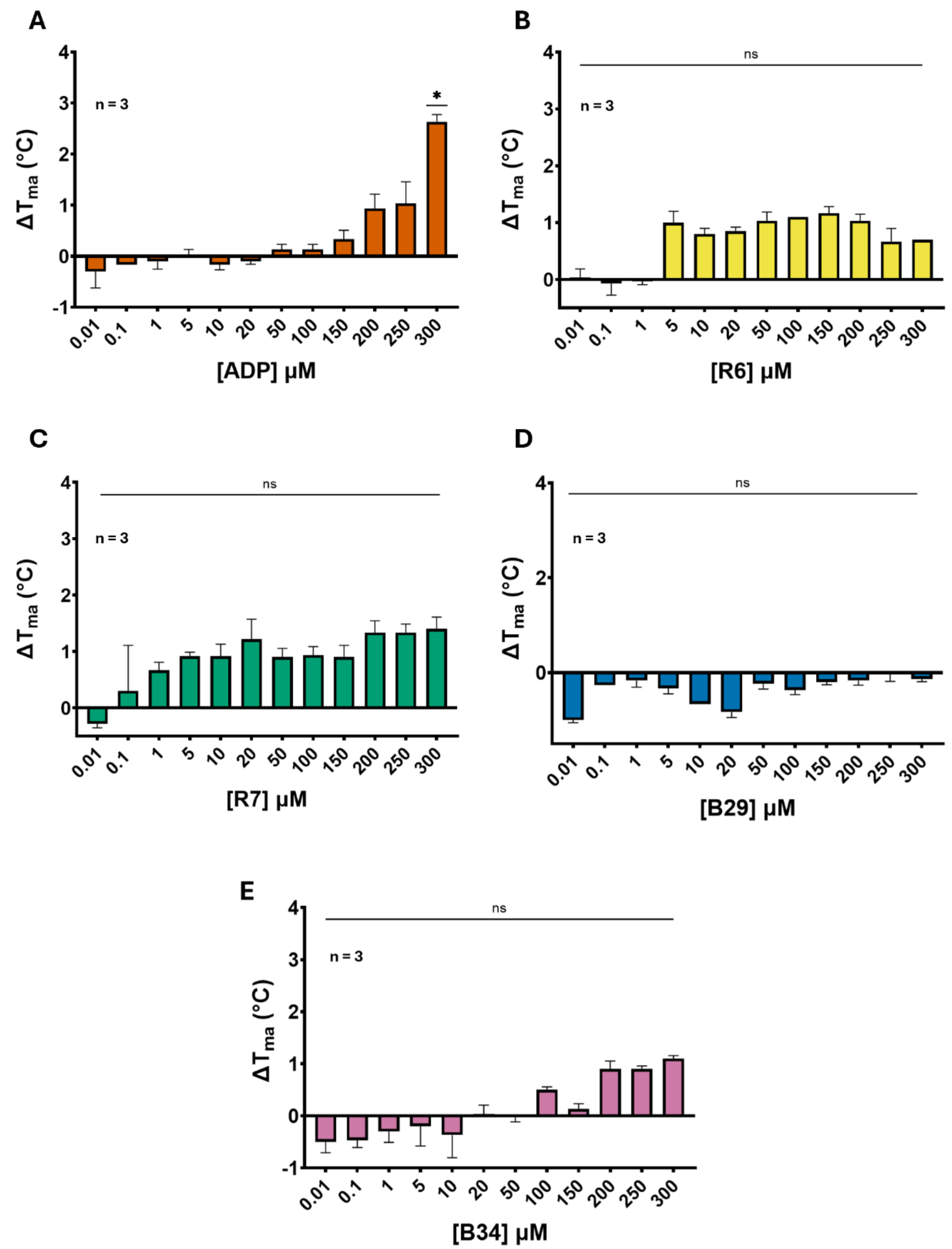


**Figure S4**. Change in melting temperature for each compound in $PhoQ_{Cyt.}$ (A-E) Change in melting temperature ($T_m$) over a range of compound concentrations. Significance determined using unpaired t-test comparisons between the $T_m$ of samples compared to the DMSO control.

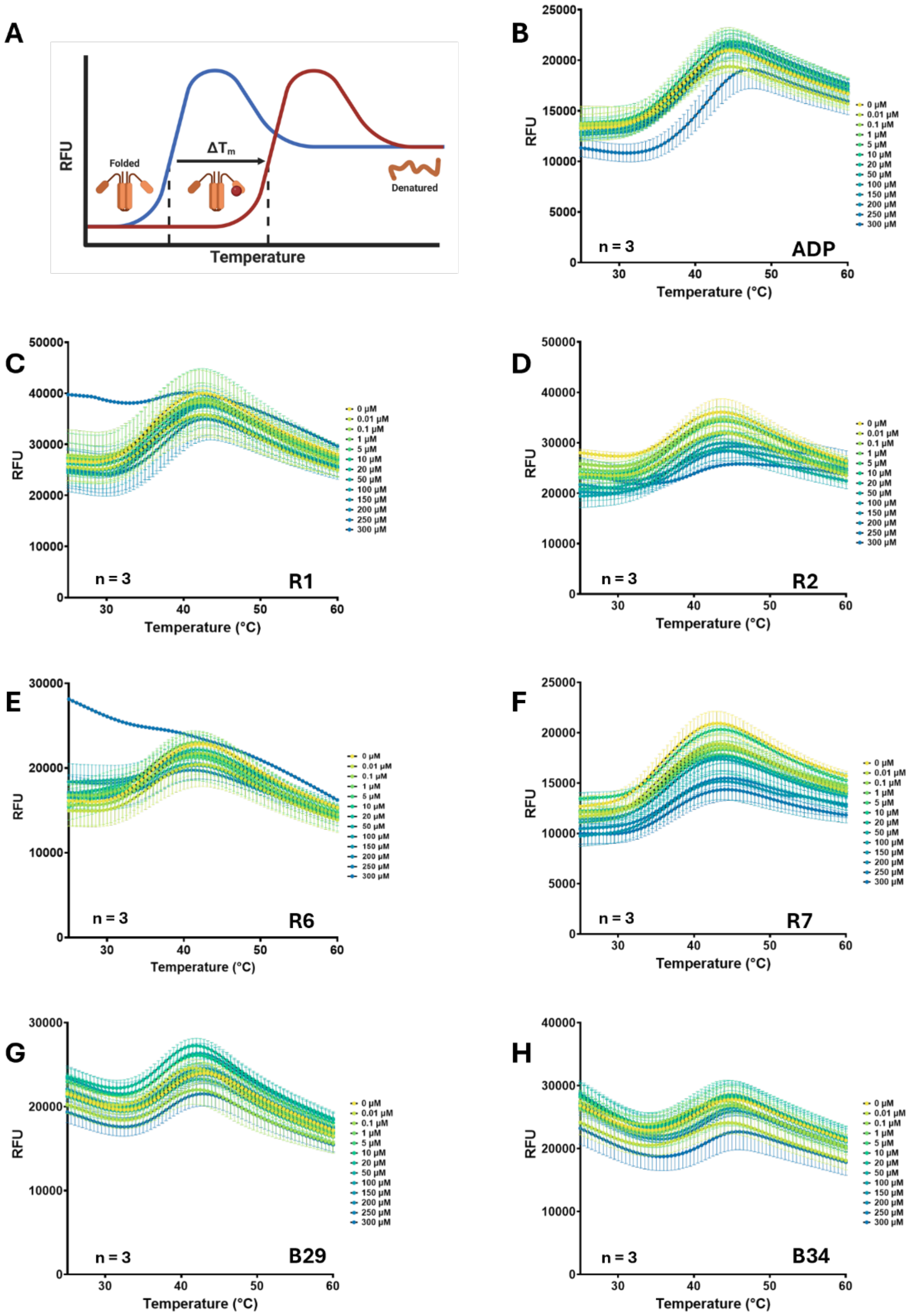


**Figure S5**. (A) Scheme depicting the method DSF uses to determine binding. (B-H). RFU curves of tested compounds in DSF with $PhoQ_{Cyt}$.
